# Spatiotemporal dynamics and stability of multi-kingdom microbial communities in hospital sinks acting as persistent pathogen reservoirs

**DOI:** 10.64898/2026.09.25.754391

**Authors:** Iván Linares-Ambohades, Natalia Guerra-Pinto, Sandra Mingo-Ramírez, Silvia Serrano-Calleja, Francisco Amaro, Ana Alastruey-Izquierdo, María Cruz Soriano, Raúl de Pablo, Val F. Lanza, Rafael Cantón, Fernando Baquero, Teresa M. Coque, Ana Elena Pérez-Cobas

## Abstract

Hospital sink drains act as persistent reservoirs for nosocomial pathogens, yet the structure and long-term stability of their associated microbial communities remain poorly known. Here, we present an extensive characterization integrating multi-kingdom meta-taxonomic profiling (16S rRNA, ITS, and 18S rRNA, n=93 samples) with culturomics (n>4000 isolates) and phenotypic resistance testing across 14 sinks from a former intensive care unit (ICU). Over a 17-month study period, we tracked community assembly and stability through the renovation and transition from an unpopulated baseline to active clinical use, encompassing both structural remodeling and functional conversion. We showed that former-ICU drains harbor a highly conserved multi-kingdom core that remains resilient despite significant structural changes and periods of inactivity. While a persistent core baseline exists, community composition changes along a continuous temporal gradient driven by structural and anthropogenic shifts. Furthermore, individual sinks assemble as unique microecosystems shaped by local environmental factors. Topological network modeling shows these communities are not random assemblages but integrated ecological units, with clinically relevant, multidrug-resistant WHO-priority pathogens deeply embedded in the network alongside often-overlooked fungi and protozoa. Together, these findings identify hospital sink drains as persistent and structured multi-kingdom microbial reservoirs that harbor clinically relevant multidrug-resistant organisms (MDROs). Their temporal stability and strong sink-specific signatures highlight the need to consider the broader microbial community when monitoring hospital environmental reservoirs and developing infection-control strategies.

**Graphical abstract:** 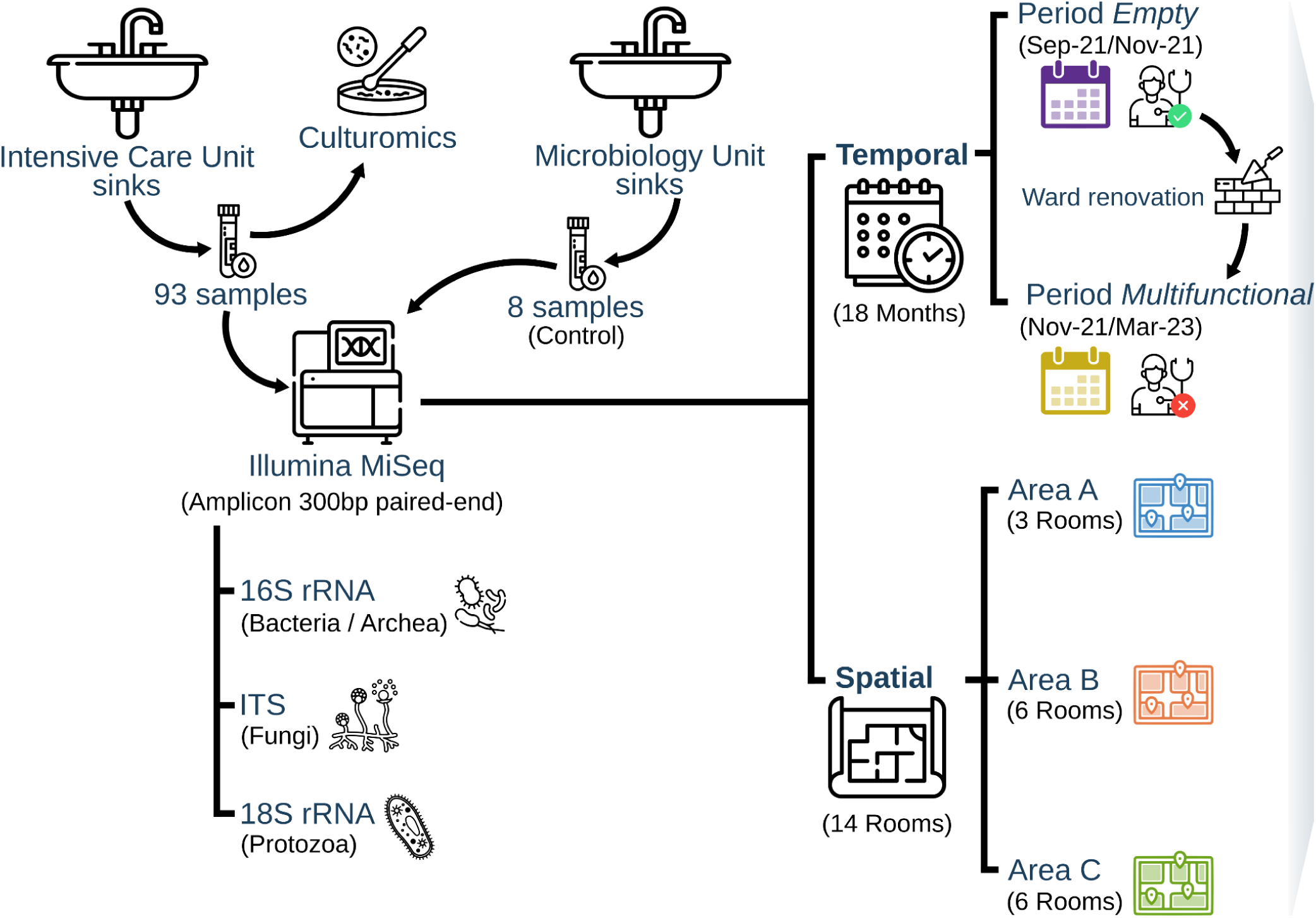

## 1. Introduction

Despite advances in diagnosis, treatment, and infection prevention, hospital-acquired infections (HAIs) remain a primary threat to global patient safety, affecting 5–7% of acute-care patients in high-income countries and up to 10% in low- and middle-income regions (Abalkhail and Marzouk, 2025). Although the World Health Organization estimates that at least 50% of HAIs are preventable through effective infection prevention and control (IPC) measures, implementation remains inconsistent across healthcare systems (“Global report on infection prevention and control 2024,” n.d.). The burden of HAIs is further exacerbated by antimicrobial resistance (AMR), which compromises treatment efficacy and increases morbidity, mortality, and healthcare costs (Coque et al., 2023; Sartelli et al., 2024). HAIs promote antibiotic use that selects for resistant bacteria, while infections caused by multidrug-resistant organisms (MDROs) require prolonged and more complex therapies, creating a self-reinforcing cycle that contributes to the global AMR crisis (Barrasa-Villar et al., 2017; Sartelli et al., 2024).

Among hospital wards, intensive care units (ICUs) experience the highest incidence of HAIs because critically ill patients are exposed to multiple risk factors, including invasive devices, mechanical ventilation, immunosuppression, and sustained broad-spectrum antimicrobial use. Consequently, approximately 20–25% of ICU patients develop an HAI during hospitalization (Vincent et al., 2020), predominantly device-associated infections such as ventilator-associated pneumonia, which are associated with mortality rates exceeding 30% (Rosenthal et al., 2024).

Increasing evidence indicates that the hospital built environment is not merely a passive recipient of microbial contamination but an active vehicle for transmission. High-touch surfaces and fomites, medical equipment, textiles, and water-associated infrastructures can serve as long-term microbial reservoirs (Russotto et al., 2017). Molecular epidemiological studies have repeatedly linked patient infections to genetically indistinguishable MDRO strains recovered from the surrounding environment (Kuczewski et al., 2022). The persistence of these organisms is largely facilitated by biofilm formation on both dry and wet interfaces where microorganisms exhibit enhanced tolerance to disinfectants and antimicrobials, facilitating long-term microbial survival and recurrent opportunities for patient colonization and infection (Birgand et al., 2022).

Within the hospital water system, plumbing infrastructure, and specifically sink drains, serve as important reservoirs of opportunistic pathogens including carbapenem-resistant Enterobacterales and *Pseudomonas aeruginosa* capable of transmission to patients (Kizny Gordon et al., 2017; Perkins et al., 2019). Their importance is especially evident in ICUs, where environmental persistence may contribute to sustained endemic transmission. Consistent with this, our previous work identified ICU sinks as long-term reservoirs of carbapenemase-producing members of the *Serratia marcescens complex*, supporting prolonged environmental persistence, plasmid dissemination, and recurrent outbreak dynamics (Aracil-Gisbert et al., 2024; “ISPB 2026 | Programme point - Hospital Water Systems Drive the Maintenance and Evolution of Acquired Antimicrobail Resistance via Robust Plasmid and Clonal Networks,” n.d.).

Recent studies have revealed that ICU sink drains harbor complex bacterial microbial communities comprising both environmental and human-associated species (Hennebique et al., 2025; Weinberger et al., 2025). However, the contribution of non-bacterial microorganisms to the ecosystem remains poorly understood. Here, we present a longitudinal multi-kingdom characterization of microbial communities inhabiting 14 ICU sink drains, integrating bacteria, archaea, fungi, and microeukaryotic profiling. Specifically, we: (i) map the intra- and inter-kingdom co-occurrence networks associated with community structure and stability; (ii) track the ecological integration of WHO-priority MDROs; and (iii) exploit a natural experiment created by the structural remodeling and functional repurposing of an ICU ward to disentangle the relative contributions of spatial and anthropogenic drivers to microbial community assembly in hospital plumbing systems.

## 2. MATERIAL AND METHODS

### 2.1. Study setting

This longitudinal study (2021–2023) was conducted at the Ramón y Cajal University Hospital, a 1,155-bed tertiary-care hospital in Madrid, Spain. We analyzed 93 sink drain (p-trap) samples collected across seven sampling time points (months 0, 2, 5, 7, 9, 13, and 17) from 14 sinks located in a decommissioned Intensive Care Unit (ICU) ward with a history of multiple outbreaks. Throughout the entire 17-month study, no ICU patients were present in the unit. Therefore, to accurately reflect the operational status of the ward, the sinks are hereafter referred to as “former-ICU sinks”. The 14 sinks were situated in rooms divided into three zones based on their former clinical layout: Area A (three former isolation rooms), Area B (six former standard ICU rooms), and Area C (five former standard ICU rooms) (**Figure 1A**). Areas A and B were directly connected through a central nursing station, whereas Area C was separated from Area B by a staff rest area, resulting in greater spatial isolation.

**Figure 1.**
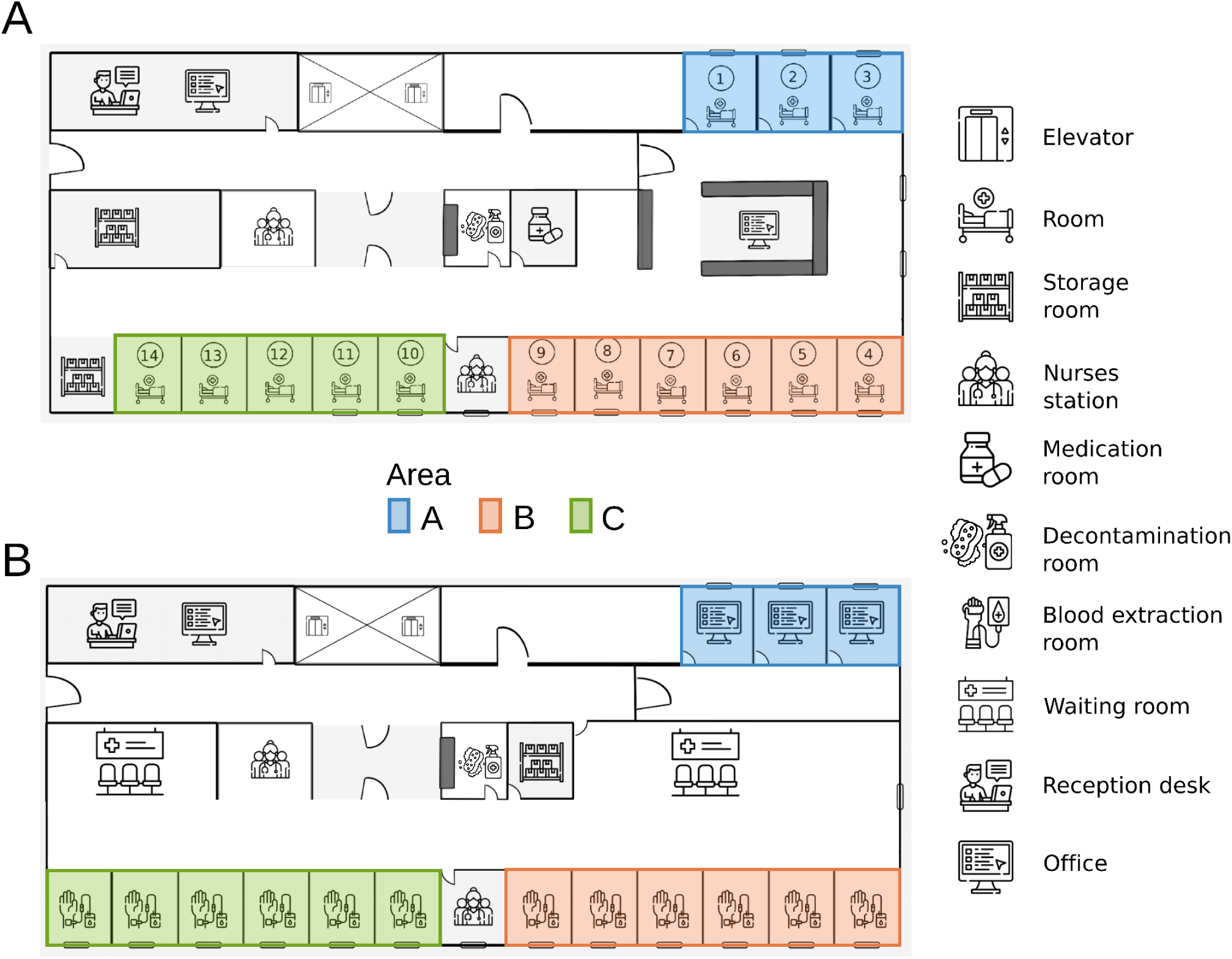
Layout of the ICU before and after remodeling. **(A)** Original ICU configuration showing the 14 sink-equipped rooms grouped into three areas (A–C) based on spatial proximity and clinical function; the study started immediately after patients’ transfer to a new ICU. **(B)** Layout after ICU remodeling and functional repurposing of the unit into a multifunctional clinical support area (offices, blood extraction, waiting, and staff areas). Sink locations and plumbing infrastructure remain unchanged while their patterns of use were substantially altered.

Prior to the initiation of sampling, all ICU clinical activity had ceased, and patients had been transferred to a newly constructed ICU facility elsewhere in the hospital. The study commenced with the original unit completely closed and unoccupied. The first 25 samples (months 0 and 2) were collected during this decommissioned baseline period, the empty-post ICU service (hereafter referred to as the “empty” phase). Following a two-month structural renovation, the ward was reopened not as an ICU, but permanently repurposed as a “multifunctional” clinical support area (without inpatient beds), comprising administrative offices, outpatient blood-drawing stations, staff facilities, and patient waiting areas. During this repurposed phase, 68 samples were collected from the same 14 sinks across months 5 to 17. Specifically, Area A was converted into offices, while Areas B and C became outpatient blood-drawing stations (**Figure 1B**).

Although the underlying plumbing infrastructure remained unchanged, sink usage shifted from total inactivity during the closure to non-inpatient clinical and administrative use. This provided a unique natural experiment to evaluate the influence of human activity on microbial community assembly in hospital plumbing systems in the complete absence of critically ill patients. Additionally, 8 sink p-trap samples were collected from the diagnostic laboratory of the Clinical Microbiology Department to serve as a comparative control group.

### 2.2. Sampling method and total DNA extraction

Water retained in each sink p-trap was homogenized by aspirating and dispensing the contents four times with a 50 mL sterile syringe. A 50 mL aliquot was then collected and centrifuged at 3000 × g for 15 minutes at 4°C. After discarding the supernatant, the resulting 2–3 mL pellet was resuspended, and a 200 μl aliquot was used for total DNA extraction using the DNeasy PowerSoil Pro Kit (QIAGEN, Germany) following the manufacturer’s protocol. DNA concentration was quantified via a Qubit® 2.0 Fluorometer (Life Technologies, Waltham, MA, USA). Negative kit controls were added but it did not produce enough DNA contaminants for sequencing in the library preparation.

### 2.3. Illumina sequencing of 16S rRNA, 18S rRNA genes, and ITS

Taxonomic profiling was performed via amplicon sequencing of three target regions using Illumina-standard adapters and primers for bacteria (16S rRNA V3-V4): 175F (5′-TCGTCGGCAGCGTCAGATGTGTATAAGAGACAGCCTACGGGNGGCWGCAG-3’) and 512R (5’-GTCTCGTGGGCTCGGAGATGTGTATAAGAGACAGGACTACHVGGGTATCTAATCC-3’) (Klindworth et al., 2013), microeukaryotic (18SrRNA V4): EUKAF (5′-GCCGCGGTAATTCCAGCTC-3’) and EUKAR (5′-CYTTCGYYCTTGATTRA-3’) (Moreno et al., 2018), and fungi (internal transcribed spacer (ITS) region): ITS1 (5’-CTTGGTCATTTAGAGGAAGTAA-3’) and ITS2 (5’-TACTTCCTCTAAATGACCAAG-3’) (Monard et al., 2013). Libraries were sequenced on an Illumina MiSeq 300 bp paired-end at the Translational Genomics (NGS) and Bioinformatics unit of Ramón y Cajal Hospital (https://www.irycis.org/en/services/11/translational-genomics-ngs-and-bioinformatics).

### 2.4. Marker gene data processing, ecological and statistical analyses

#### Reads processing and taxonomic assignment

Raw paired-end FASTQ reads were quality-assessed with FastQC (v0.12.1), removed of adapters and primers using Cutadapt v5.0 (Martin, 2011), and quality-filtered via Trimmomatic v5.0 (Bolger et al., 2014). Potential host contamination was eliminated by filtering reads that aligned to the human reference genome using Bowtie (v1.3.1) (Langmead and Salzberg, 2012).

Taxonomic classification was performed with Kraken2 v1.3.1 (Wood et al., 2019) using marker-specific reference databases mapped to the NCBI taxonomy. For prokaryotic profiling, a precompiled 16S reference dataset from the Ribosomal Database Project (RDP) release 11.5 (Cole et al., 2014) was used. For eukaryotic small-subunit (18SrRNA) profiling, a custom database was constructed by integrating EukRibo v2 (Berney, 2022), PR2 SSU release v5.0.0 (Bass et al., 2013), a 2024 SILVA package (Quast et al., 2013), and the RefSeq “protozoa” library. For fungal ITS profiling, a custom fungal ITS database was assembled by merging the UNITE dynamic release 04.04.2024 (Abarenkov et al., 2024) with the RefSeq “fungi” library. Both custom databases were compiled following standard Kraken2 build instructions. The amplicon custom analyses are freely available on GitHub (https://github.com/ivan24n/sink_micro_analysis).

#### Core determination and stability

Marker-specific relative abundance tables (16S rRNA, 18S rRNA, and ITS) were processed in R to define the core microbiome (≥80% sample prevalence, and 70% for the 18S rRNA) and determine mean relative abundances. Longitudinal dynamics for the ten most abundant core genera were evaluated using (Shields-Cutler et al., 2018). Specifically for the 16S rRNA, we used the trendyspliner function to test for significant non-zero temporal trends by comparing smoothing splines against permutation-generated null distributions, while the permuspliner function facilitated pairwise comparisons between study areas, utilizing false discovery rate adjustments to detect significant differences in taxon abundances between groups throughout the study period.

#### Microbial associations prediction

To evaluate ecological associations, genus-level networks were constructed for each marker (16S rRNA, 18S rRNA, and ITS) using taxa with >10% sample prevalence and >0.01% total relative abundance. Pairwise co-occurrences were estimated via Maximum Likelihood Estimation (MLE) via the CooccurrenceAffinity R package (v1.02), retaining only significant (p<0.05), positive αMLE interactions.

Networks were visualized and analyzed in Gephi v0.10.1. Global topological metrics (average degree, diameter, density, connected components, modularity, and average path length) were computed and validated against random null models matched for node count and edge density. Genera were organized into distinct clusters using modularity-based algorithms to identify highly interconnected ecological units. Following modularity-based clustering, nodes were categorized by phylum. The 16S rRNA network was cross-referenced with the 2024 WHO Bacterial Priority Pathogens List to assess the centrality of high-risk taxa. Final edges were weighted by their corresponding αMLE affinity scores.

#### Alpha and beta diversity

Sample rarefaction was performed using the vegan R package (v2.7-1) based on genus-level diversity curves to ensure standardized comparisons. Alpha diversity (Shannon index, Chao1 richness estimator, and Berger-Parker index metrics) was evaluated across groups using Bonferroni-corrected Wilcoxon rank-sum tests.

Beta diversity was visualized via Non-Metric Multidimensional Scaling (NMDS) of Bray-Curtis and Jaccard distance matrices. Global compositional variance was tested via PERMANOVA (adonis2 function), with spatial pairwise comparisons (Areas A, B, and C) performed through pairwise permutation tests using the pairwiseAdonis package (v0.4.1). All diversity and compositional analyses were executed using a suite of R packages, including phyloseq (v1.52.0), microbiome (v1.30.0), and ggplot2 (v4.0.0) for high-quality data visualization. Taxon overlap between groups was mapped with the ggvenn R package (v0.1.19). and differential abundance was tested via Analysis of Compositions of Microbiomes with Bias Correction 2 (ANCOM-BC2 v2.10.1) (Lin and Peddada, 2024).

#### Self-Organizing Map analysis of temporal microbial profiles

To cluster taxa with shared temporal trajectories while preserving the data’s continuous structure, Self-Organizing Maps (SOMs) were applied. Prior to mapping, the dataset was filtered to retain taxa present in at least three time points. Counts underwent count-zero multiplicative (CZM) imputation (zCompositions v1.6.0), centered log-ratio (CLR) transformation (compositions v2.0-9). Rows were scaled to emphasize the shape of the temporal profile.

A rectangular SOM grid was constructed with a size approximately 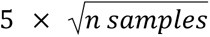 to balance profile resolution and representation (Tian et al., 2014). The SOM was trained for 200 iterations with a linearly decreasing learning rate from 0.05 to 0.01, using Euclidean distance, implemented with the R Kohonen package v3.0.12. Model quality was evaluated using quantization error (the average distance of data points to their best-matching unit) and topographic error (the fraction of data points whose two closest units are not adjacent), averaged over 50 independent runs. Node codes from the trained SOM were clustered using Ward’s hierarchical method (cluster v2.1.6, factoextra v1.0.7). The optimal number of clusters was determined by maximizing the mean silhouette width computed on taxa assignments. Cluster stability was assessed across runs using the Adjusted Rand Index (mclust v6.1.2). Temporal dynamics of clusters were visualized as median CLR profiles for each cluster, enabling the identification of recurring temporal patterns.

### 2.5. Bacterial identification, antimicrobial susceptibility testing, and resistance gene detection

Undiluted 100 µl aliquots of vortex-homogenized sink water were spread-plated and incubated aerobically at 37°C for 24h across three CHROMagar™ media: ESBL Biplate (for extended-spectrum β-lactamase-producing Gram-negative bacteria), mSuperCARBA™ (for carbapenemase-resistant *Enterobacterales*), and Orientation (for general isolation and differentiation). Representative colonies displaying distinct morphotypes were identified by matrix-assisted laser desorption/ionization time-of-flight mass spectrometry (MALDI-TOF MS, MALDI Biotyper, Bruker; Germany), accepting identifications with a score >2.0. Subcultured isolates were stored at −80°C in Luria-Bertani (LB) broth supplemented with 15% (v/v) glycerol.

Disk-diffusion antimicrobial susceptibility testing was performed on Mueller-Hinton agar using 0.5 McFarland suspensions of clinically relevant isolates. Antibiotic disks (Bio-Rad) were applied using a 7-disk dispenser and incubated at 37 °C for 24 h. Inhibition zone diameters were interpreted according to the European Committee on Antimicrobial Susceptibility Testing (EUCAST) clinical breakpoints and epidemiological cut-off values (ECOFFs) (https://www.eucast.org/bacteria/). For Gram-negative isolates (Chryseobacteria, Enterobacteriaceae, and *Pseudomonas*) susceptibility testing included ampicillin (10µg), amoxicillin-clavulanic acid (20:10µg), cefoxitin (30µg), ceftazidime (10µg), cefotaxime (5µg), cefepime (30µg), aztreonam (30µg), temocillin (30µg), and meropenem (10µg). Gram-positive isolates were tested against vancomycin (5 µg), streptomycin (500 µg), gentamicin (500 µg), and erythromycin (15 µg). Extended-spectrum β-lactamase (ESBL) and carbapenemase genes were detected by multiplex PCR as previously described (Monstein et al., 2007; Poirel et al., 2011).

## 3. Results

### 3.1. Persistent multi-kingdom core microbiome defines the microbial signature of ICU sink drains

After applying quality filtering and human-read removal usingBowtie2, the final dataset comprised 93 bacterial (16S), 34 fungal (ITS), and 57 microeukaryotic (18S) samples. 1Median sequencing depths were 110591 reads (IQR: 89814–145283), 120446 reads (IQR: 90683–151739), and 83632 reads (IQR: 58689–124395) for the 16S, ITS, and 18S datasets, respectively (**Supplementary Table 1**).

To identify key members contributing to long-term community persistence, we defined the core microbiome as genera detected in >80% of samples for each marker (**Supplementary Table 2**). The bacterial core comprised 92 genera accounting for an average of 76.97% of the total community relative abundance, and was dominated by members of the phyla Pseudomonadota and Bacteroidota. Fifteen genera were found in 100% of the samples, including the highly abundant *Pseudomonas*, *Comamonas*, *Burkholderia*, *Sphingomonas*, and *Sideroxydans*, all from Pseudomonadota (**Figure 2A**). Other abundant and highly prevalent members included, such as *Magnetospirillum* (Pseudomonadota) together with *Cloacibacterium*, *Elizabethkingia*, and *Flavobacterium* (Bacteroidota) (**Figure 2A**).

**Figure 2.**
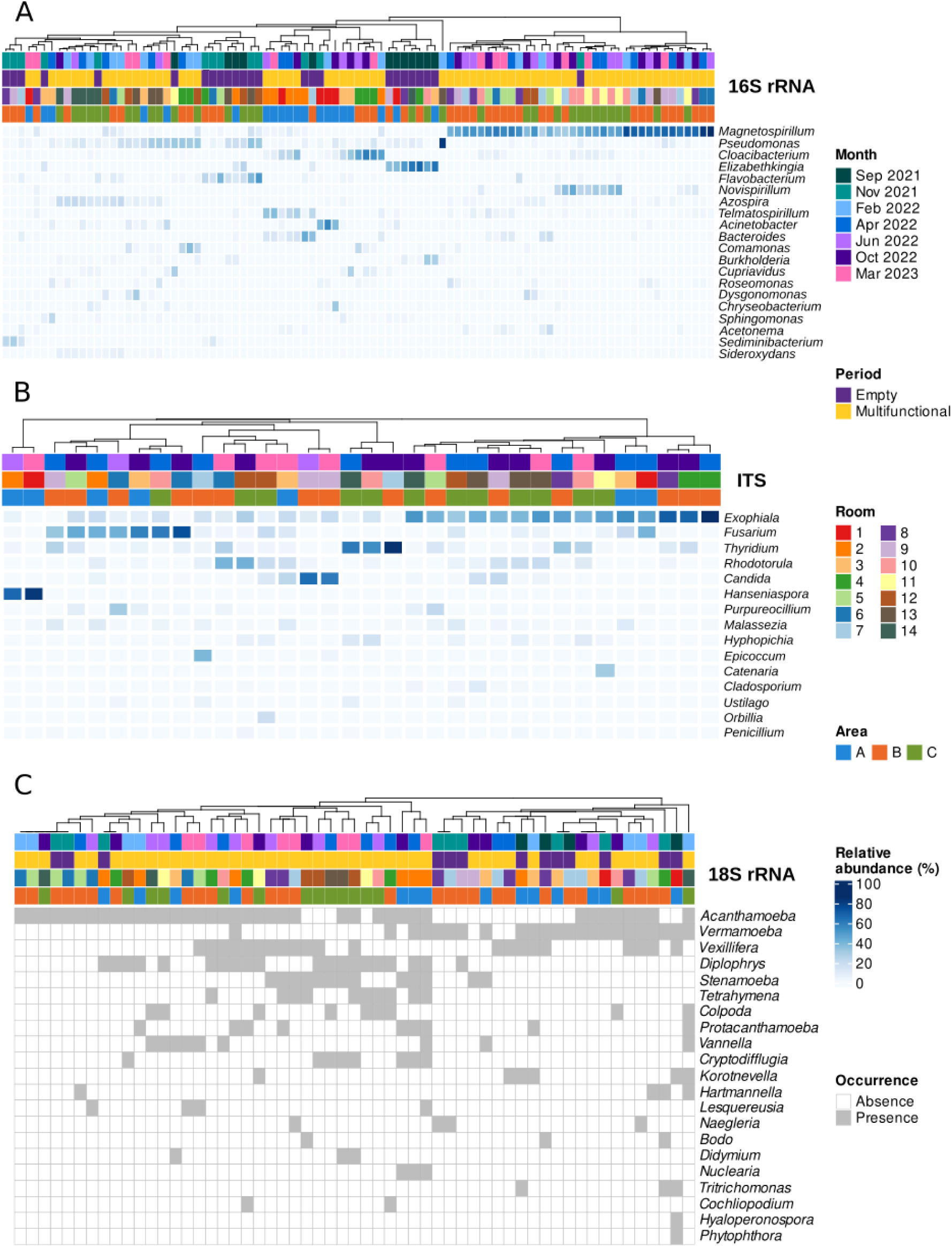
Taxonomic clustering heatmaps of the ICU sink cohort. **(A)** Relative abundance (%) of dominant bacterial taxa (16S rRNA, *n=*93). **(B)** Relative abundance (%) of dominant fungal taxa (ITS, *n=*34). **(C)** Occurrence of the most prevalent microeukaryotes (18S rRNA, *n=*57). Columns are annotated by sample room, space area, sampling month, and study period.

The fungal core was markedly smaller, comprising only 8 genera that collectively represented 81.18% of the fungal community. It was dominated by members of the Ascomycota (*Exophiala*, *Fusarium*, *Thyridium*, *Candida*) and Basidiomycota (*Rhodotorula*, *Malassezia*) (**Figure 2B**). In contrast, the microeukaryotic community exhibited limited taxonomic persistence. Only the amoebozoan genus *Acanthamoeba* surpassed the 70% prevalence, while all remaining taxa remained well below this threshold **(Figure 2C)**.

To determine whether former-ICU sink drains harbored specific microbial signatures, we compared their communities with those of sink p-traps from the hospital Clinical Microbiology laboratory (**Supplementary Figure 1**). Bacterial communities in former-ICU sinks exhibited significantly higher richness than those from the Clinical Microbiology laboratory (Chao1 richness estimator, *p*<0.001), whereas community evenness and dominance did not differ significantly (Shannon *p*=0.1, Berger-Parker *p*=0.1; **Supplementary Figure 1A**). Conversely, the beta-diversity analysis revealed significant differences in composition between units (Bray-Curtis, R² = 0.045, *p* = 0.001; Jaccard index R² = 0.05, *p* = 0.001; **Supplementary Figure 1B**). Despite 347 shared genera, each unit displayed a distinct taxonomic profile (**Supplementary Figure 1C**).

The mycobiome showed a similar pattern (**Supplementary Figure 1D**); while richness differed significantly between units (Chao1 estimator, *p*=0.011), no significant differences were observed in Shannon (*p*=0.128) or Berger-Parker (*p*=0.063) diversity indices. Nevertheless, beta-diversity analysis showed unique fungal profiles despite sharing 132 genera (Bray-Curtis, R^2^=0.082, *p*=0.002; Jaccard, R^2^=0.059, *p*=0.001; **Supplementary Figure 1E-F**).

Finally, microeukaryotic communities followed the same overall trend. Former-ICU sinks harbored significantly greater richness than the control sinks (observed species, *p*=0.03; **Supplementary Figure 1G**), while taxonomic overlap between the two units was low, with only three shared genera representing 13% of the total generic richness (**Supplementary Figure 1H**). Most microeukaryotic taxa detected in former-ICU sinks were absent in the control samples (**Supplementary Figure 1I**).

### 3.2. Sink-specific microbial patterns of a hospital unit

Despite a unit-wide core microbiota, bacterial communities exhibited strong sink-specific signatures. Both abundance-based (Bray–Curtis) and incidence-based (Jaccard) distances were significantly greater between different sinks than within the same sink (Wilcoxon test, p<0.001; **Figure 3A**), consistent with PERMANOVA results (Bray-Curtis R^2^=0.057, p<0.001; Jaccard R^2^=0.033, p<0.001; **Figure 3B**). Across the 91 pairwise comparisons among the 14 sinks (Pairwise Adonis), 65 Bray-Curtis and 74 Jaccard comparisons were significant (*p*<0.05), with sinks 1, 2, 3, 4, 13, and 14 showing the greatest divergence from the rest of the unit (**Supplementary Table 3**).

**Figure 3.**
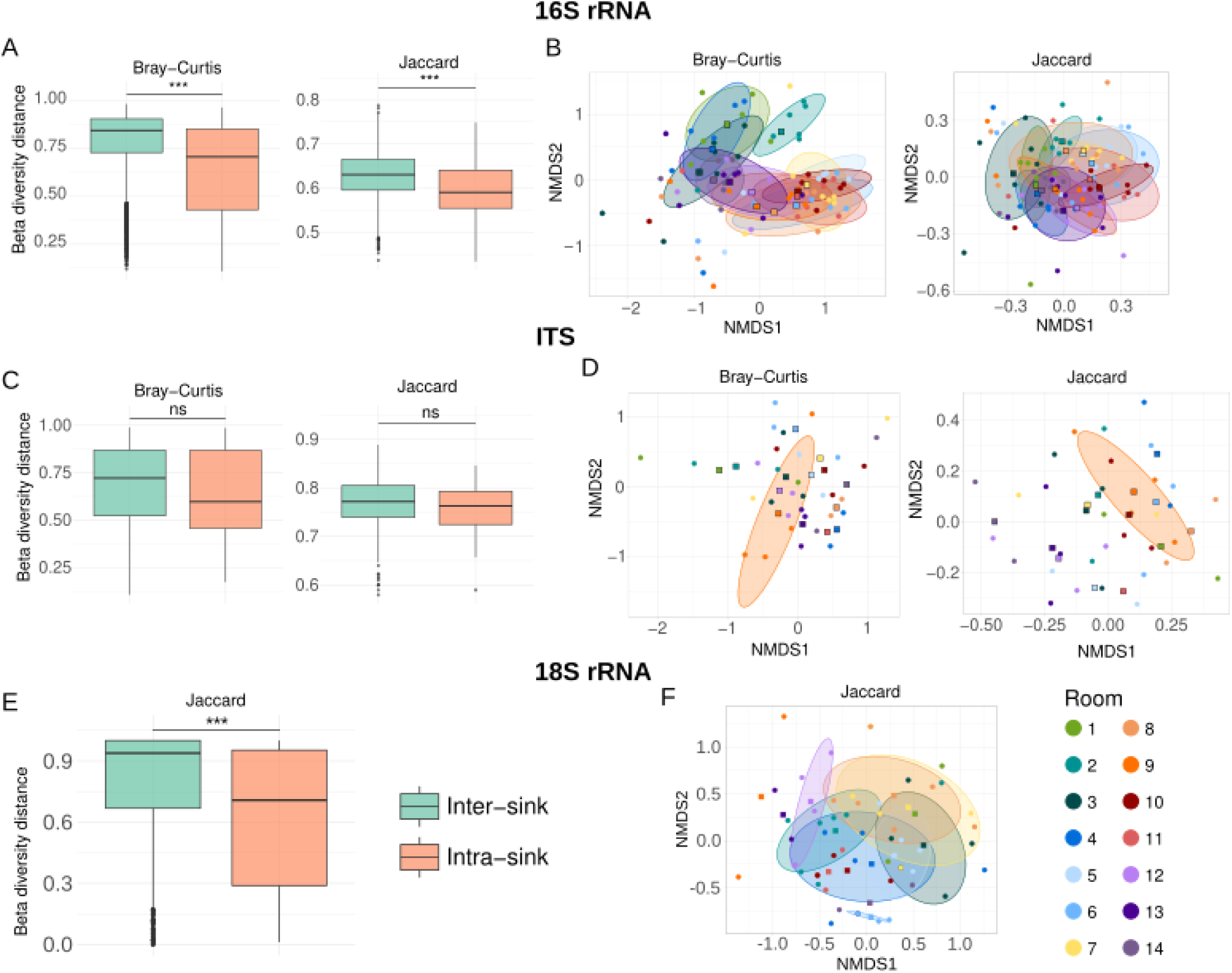
Beta diversity comparison of sink microbial communities. Inter-versus intra-sink distance boxplots and corresponding Non-Metric Multidimensional Scaling (NMDS) plots for: **(A, B)** bacteria (16S rRNA, *n=*93; Bray-Curtis and Jaccard); **(C, D)** fungi (ITS, *n=*34; Bray-Curtis and Jaccard); and **(E, F)** microeukaryotes (18S rRNA, *n=*57; Jaccard). Boxplots show inter- and intra-sink distances compared (**A, C, E**) using Bonferroni-corrected Wilcoxon tests: *p < 0.05, **p < 0.01, ***p < 0.001. “ns“: non-significant differences (p>0.05).

Fungal communities showed a weaker spatial structure. Inter-sink distances tended to exceed intra-sink distances but did not reach statistical significance (Wilcoxon p>0.05; **Figure 3C**). Nonetheless, PERMANOVA detected statistically significant differences in community composition among sinks (Bray-Curtis R^2^=0.066, p<0.05; Jaccard R^2^=0.040, p<0.05; **Figure 3D**), although no individual sink pairs remained significant in subsequent pairwise comparisons (**Supplementary Table 3**). This result likely reflects the diminished statistical power of the smaller ITS dataset (*n=*34) rather than a true absence of spatial structuring.

Microeukaryotic communities also displayed spatial heterogeneity. Jaccard distances were significantly greater between sinks than within sinks (Wilcoxon *p*<0.001; **Figure 3E**), and PERMANOVA confirmed a significant sink-specific effect on community composition (R^2^=0.04, *p*<0.05; **Figure 3F**). However, pairwise analysis identified only a sparse subset of significant differences, primarily involving sinks 11 and 14 (**Supplementary Table 3**). Collectively, these multi-kingdom analysis indicate that individual sinks harbor distinct microbial assemblages that persist despite the presence of a shared ICU-wide core microbiota. Although sink identity explained only a modest proportion of the overall community variation, the consistent spatial signal across microbial kingdoms supports the existence of localized ecological niches within the hospital water network.

### 3.3. Multi-kingdom microbial co-occurrence analyses

To explore possible ecological associations, we constructed intra-kingdom (**Supplementary Figure 2-4**) and inter-kingdom (**Figure 4**) co-occurrence networks. Separate inter-kingdom networks were constructed because integrating all three kingdoms would have reduced the analysis to the subset of samples with complete multi-marker data and increased sparsity due to the markedly different prevalence of fungal and microeukaryotic taxa, limiting network resolution.

**Figure 4.**
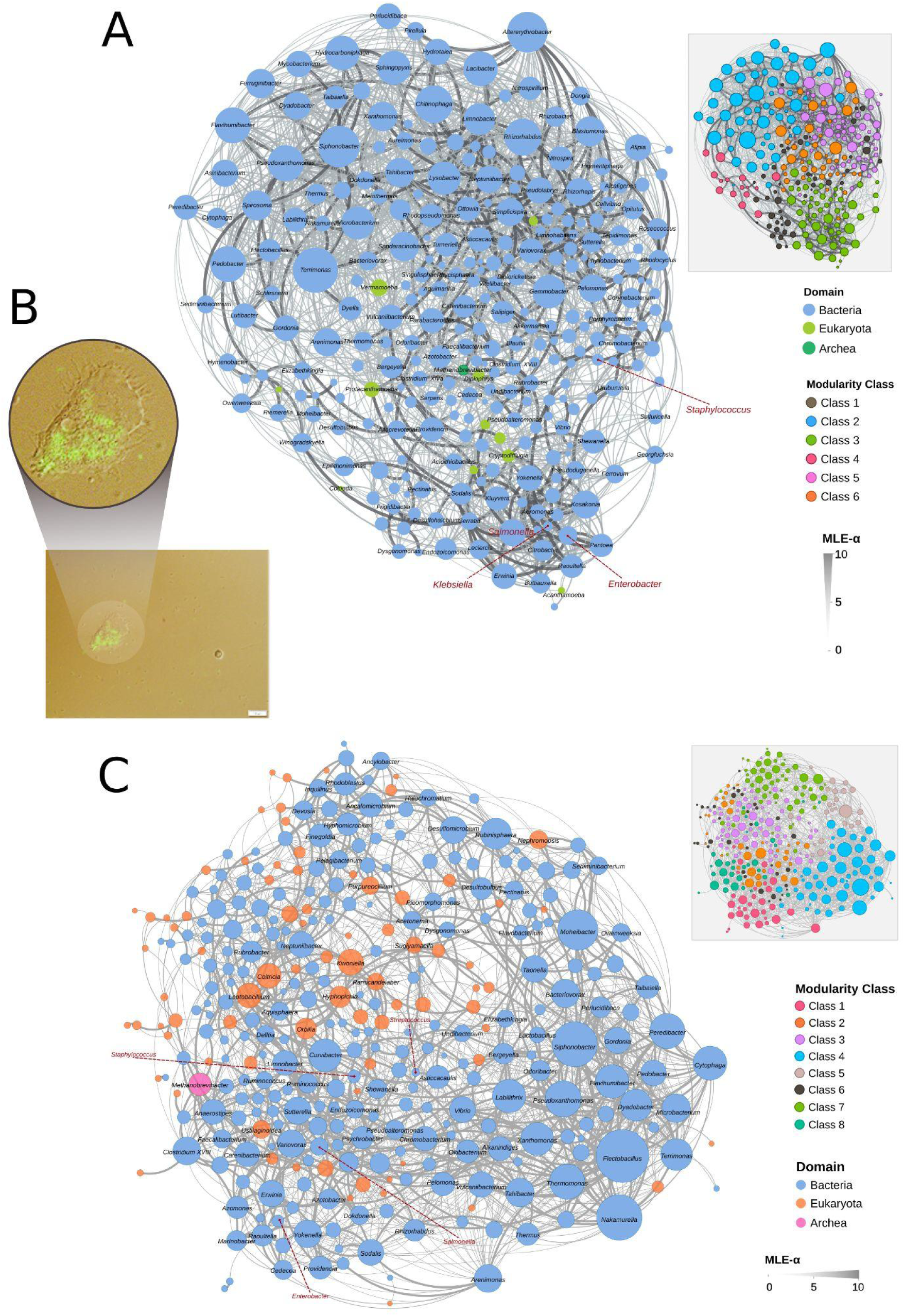
Inter-kingdom microbial co-occurrence networks. **(A)** Prokaryote–microeukaryote network with nodes colored by domain. The inset (bottom-left) shows the six network modules identified by modularity analysis (modules 1–6, n=48, 55, 57, 16, 61 and 40 nodes, respectively). **(B)** Fluorescence microscopy image of *Acanthamoeba* sp. isolate B2RYC ingesting GFP-expressing *Burkholderia cenocepacia* and unlabeled indigenous bacteria. The image represents merged Nomarski and fluorescence microscopy. White scale bar represents 10 µm. **(C)** Prokaryote–fungi network with nodes colored by domain (Bacteria, Archaea, Fungi). The inset (top-right) shows the 8 network modules (modules1–8, n=31, 34, 66, 53, 23, 35, 56, and 31). Red nodes denote WHO priority pathogens. Main network panels display high-confidence strong associations (αMLE>5, p<0.05), while modularity insets retain all significant associations (p<0.05) regardless of αMLE, to visualize the total global topology. Modularity class is a group number assigned to nodes by a community-detection algorithm. αMLE is the maximum likelihood estimate of the log-odds parameter alpha (“CooccurrenceAffinity” R package).

The integrated 16S rRNA–18S rRNA network comprised 279 nodes (mean degree ≈ 15; **Figure 4A**) and exhibited a strongly modular, non-random topology (**Table 1**), partitioning into six groups (**Supplementary Table 4**). Most groups were almost exclusively bacterial (>95% bacteria), whereas two bacteria-dominated (89.5% and 94.5% bacteria, clusters 2 and 3) were bridged by highly connected microeukaryotic hubs, including *Vermamoeba*, *Protacanthamoeba*, *Stenamoeba*, *Tetrahymena*, and *Acanthamoeba*. Specifically, group 3 harbored a tightly connected assemblage of clinically relevant Enterobacterales (*Citrobacter*, *Serratia*, *Raoultella*, *Enterobacter*, *Klebsiella*) together with *Streptococcus*; group 2 hosted highly connected bacterial genera *Terrimonas*, *Siphonobacter*, *Altererythrobacter*, and *Chitinophaga*, with *Vermamoeba* and *Protacanthamoeba* representing the most connected microeukaryotic nodes in the entire network. Overall, eukaryotic taxa represented a minor fraction of the network (∼4%) and acted as key connectors of bacterial-groups rather than forming a group of their own. We showed a microscopy of a prevalent protozoan, *Acanthamoeba* isolated from one of the sinks interacting with *Burkolderia*, a taxon found in 100% of the sinks (**Figure 4B**).

**Table 1.** Topology properties of the 16S rRNA, ITS, 16S rRNA-18S rRNA, and 16S rRNA-ITS networks.

|  | 16S-18S rRNA /<br>Random | 16S rRNA-ITS /<br>Random | 16S rRNA /<br>Random | ITS / Random |
| --- | --- | --- | --- | --- |
| Nodes | 279 / 279 | 334 / 334 | 277 / 277 | 33 / 33 |
| Edges | 2098 / 2069 | 1786 / 1716 | 3284 / 3353 | 37 / 35 |
| Average degree | 42.9 / 7.42 | 10.70 / 5.14 | 23.71 / 12.11 | 2.24 / 1.06 |
| Diameter | 6 / 4 | 8 / 5 | 6 / 3 | 7 / 6 |
| Density | 0.06 / 0.05 | 0.03 / 0.03 | 0.09 / 0.09 | 0.07 / 0.07 |
| Connected<br>components | 1 / 1 | 3 / 1 | 1 / 1 | 5 / 6 |
| Modularity | 0.50 / 0.23 | 0.56 / 0.03 | 0.498 / 0.167 | 0.65 / 0.60 |
| Average path<br>length | 2.74 / 2.38 | 3.23 / 2.74 | 2.36 / 2.02 | 2.76 / 3.42 |

The integrated 16S–ITS co-occurrence network comprised 334 nodes connected by 1,786 edges) and was partitioned into nine discrete groups (**Table 1**, **Figure 4B, Supplementary Table 4**). The most densely interconnected (group 4 in blue, average degree 16.5) was dominated by bacteria (92%), and anchored by highly connected hubs such as *Pseudoxanthomonas* and *Thermomonas*. Conversely, group 6 was fungal-dominated (68%); despite lower overall connectivity, it drove specific cross-domain pairings. The largest group (66 nodes, 3), exhibited a mixed composition (62% bacteria and 38% fungi), illustrating extensive cross-kingdom connectivity. Finally, high-priority antibiotic-resistant WHO pathogens (*Enterobacter*, *Staphylococcus*, *Streptococcus*) were integrated throughout the global network rather than restricted to individual modules.

Intra-kingdom networks also exhibited modular topologies (**Table 1**). The bacterial network (6 groups; **Supplementary Figures 2-4**) retained the *Enterobacterales* enriched community identified in the integrated network whereas the fungal network (7 groups; **Supplementary Figure 3**) was comparatively sparse and fragmented, with a central Ascomycota-dominated group (3) that included the opportunistic respiratory pathogen *Aspergillus*.

### 3.4. Culturable diversity, spatial distribution, and AMR gene profiles

Over the 17-month study period, culturomics yielded 4023 bacterial isolates across 30 genera from the 93 cultured sink samples, including seven bacterial genera in the WHO priority list of antibiotic resistant pathogens (**Table 2**). While priority taxa spanned all ICU zones, their distribution was strongly spatially structured (Fisher’s Exact Test, p < 0.001); *Serratia* was recovered exclusively from Area B, and most genera displayed localized dominance, whereas *Pseudomonas* remained ubiquitous throughout the unit. Following the conversion of the ICU into a multifunctional space, the culturable community underwent a significant compositional shift (Fisher’s Exact test, ps < 0.001), accompanied by a modest increase in overall isolated recovery. Associated with this change we found that *Escherichia*, *Phytobacter*, *Citrobacter*, *Klebsiella* and *Serratia* increased, whereas *Enterobacter* and *Pseudomonas* decreased.

**Table 2.** Number of isolates per genus by area, unit use, and resistance genes harbored.

| Genus | A | B | C | T. Fisher (Area) | Empty | Multifunctional | T. Fisher (Period) | $bla_{CTXM-15}$ | $bla_{KPC-3}$ | $bla_{OXA-48}$ | $bla_{VIM-1}$ | $bla_{GES-5}$ |
| --- | --- | --- | --- | --- | --- | --- | --- | --- | --- | --- | --- | --- |
| *Klebsiella* | 87/416 (21%) | 147/917 (16%) | 52/781 (6.6%) | $X^2=55.81$<br>p.adj < 0.001 | 63/740 (8.5%) | 223/1.373 (16.2%) | $X^2=24.54$<br>p.adj < 0.001 | 7/8 (87.5%) | 3/8 (37.5%) | 0 | 0 | 0 |
| *Enterobacter* | 117/416 (28.1%) | 165/917 (83.7%) | 95/781 (12.2%) | $X^2=47.21$<br>p.adj < 0.001 | 156/740 (21.1%) | 221/1.373 (16.1%) | $X^2=8.15$<br>p.adj < 0.005 | 3/164 (1.8%) | 20/164 (12.2%) | 0 | 117/164 (71.3%) | 0 |
| *Citrobacter* | 16/416 (3.8%) | 152/917 (16.5%) | 247/781 (31.6%) | $X^2=142.34$<br>p.adj < 0.001 | 123/740 (16.6%) | 292/1.373 (21.2%) | $X^2=6.58$<br>p.adj < 0.05 | 0 | 0 | 0 | 25/44 (56.8%) | 0 |
| *Escherichia* | 19/416 (4.6%) | 8/917 (0.9%) | 2/781 (0.2%) | $X^2=40.27$<br>p.adj < 0.001 | 0/740 (0%) | 29/1.373 (2.1%) | $X^2=15.85$<br>p.adj < 0.001 | 0 | 0 | 0 | 0 | 0 |
| *Phytobacter* | 18/416 (4.3%) | 71/917 (7.7%) | 120/781 (15.4%) | $X^2=45.47$<br>p.adj < 0.001 | 38/740 (5.1%) | 171/1.373 (12.4%) | $X^2=28.90$<br>p.adj < 0.001 | 5/12 (41.6%) | 0 | 0 | 0 | 0 |
| *Serratia* | 0/416 (0%) | 20/917 (2.2%) | 0/781 (0%) | $X^2=26.36$<br>p.adj < 0.001 | 5/740 (0.6%) | 15/1.373 (1.1%) | $X^2=0.89$<br>p.adj=0.37 | 0 | 0 | 0 | 0 | 0 |
| *Pseudomonas* | 68/416 (16.3%) | 155/917 (17%) | 144/781 (18.4%) | $X^2=1.06$<br>p.adj = 0.58 | 180/740 (24.3%) | 187/1.373 (13.6%) | $X^2=38.39$<br>p.adj < 0.001 | 0 | 0 | 0 | 8/53 (15.1%) | 17/53 (32.1%) |

Genotypic characterization identified *bla*_VIM-1_ as the most prevalent resistance determinant spanning *Enterobacter*, *Citrobacter*, and *Pseudomonas*, followed by *bla*_CTXM-15_ which was restricted to *Klebsiella*. *Enterobacter* exhibited the highest diversity of acquired resistance genes (*bla*_CTXM-15_, *bla*_KPC-3_, and *bla*_VIM-1_), followed by *Klebsiella* and *Pseudomonas* (two determinants each). All remaining genera carried at most a single resistance gene.

### 3.5. Area-specific spatial structuring of ICU sink communities

Bacterial communities exhibited clear spatial structuring across the three areas. Chao1 richness was significantly elevated in Area A relative to Areas B (*p*=0.034) and C (*p*=0.024), whereas evenness and dominance remained comparable across the units (*p*>0.05; **Figure 5A, Supplementary Figure 4A**). Beta diversity analysis confirmed significant compositional differences among areas (Bray-Curtis R^2^= 0.10, p=0.001; Jaccard R^2^=0.06, *p*=0.001). Pairwise comparisons indicated that this pattern was driven primarily by the separation of Area A from Area B and Area C, while differences between Areas B and C were comparatively modest (**Figure 5B**, **Supplementary Figure 4B, Supplementary Table 3**). Despite a large shared core (785 genera, 65.9%; **Supplementary Figure 4C**), relative taxon abundances vary significantly among areas (**Figure 5C**). *Acinetobacter* predominated in Area A whereas *Magnetospirillum* was associated with Areas B and C (**Figure 5C**). ANCOM-BC2 identified 41 and 29 differentially abundant taxa between Area A and Areas B and C, respectively, compared with only 22 and10 taxa between Areas B and C (**Supplementary Table 5**). Longitudinal spline modeling further demonstrated that these spatial differences remained stable over time, with significant spatio-temporal trends driven strictly by *Magnetospirillum* (Areas A vs. B and C) and *Cloacibacterium*, *Telmatospirillum*, and *Bacteroides* (Area A vs. C) (permuspliner analysis; **Supplementary Figure 4D**).

**Figure 5.**
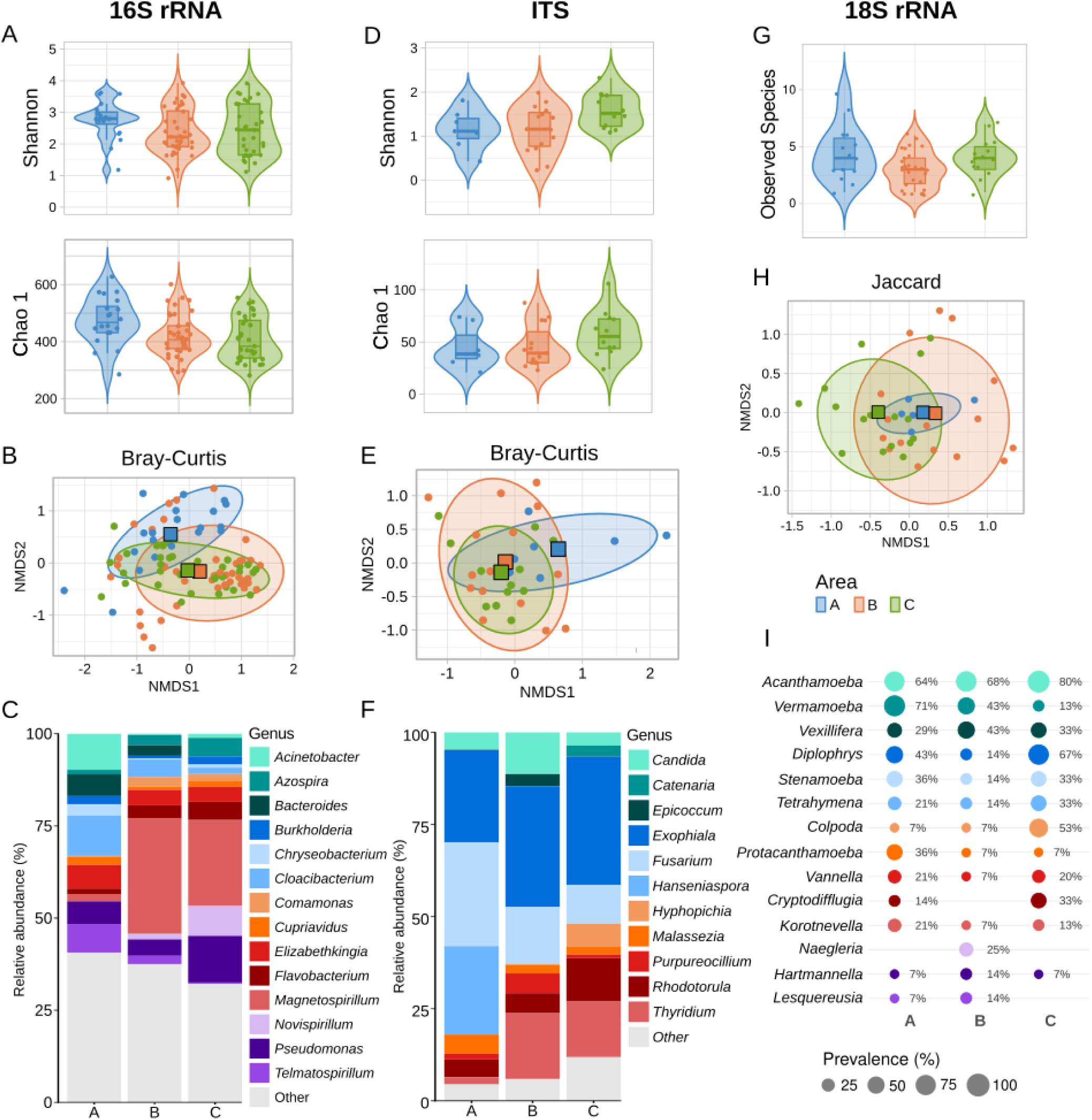
Spatial variation in microbial diversity and composition across ICU areas. Bacterial communities (16S rRNA, n=93)**: (A)** alpha-diversity metrics (Shannon index and Chao 1 richness); **(B)** Non-metric multidimensional scaling (NMDS) plot based on Bray-Curtis dissimilarities; **(C)** relative abundance of the most abundant taxa (Other < 2%). Fungal communities (**ITS,** n=34)**: (D)** alpha-diversity metrics (Shannon index and Chao 1 richness); **(E)** NMDS plot based on Bray-Curtis dissimilarities; **(F)** relative abundance (%) of the most abundant taxa (Other < 2%). Microeukaryotic communities (**18S rRNA,** n=57)**: (G)** observed richness. **(H)** NMDS ordination based on the Jaccard dissimilarities; **(I)** prevalence (%) of the most frequently detected taxa.

The mycobiome showed significantly lower richness in Area A than in Area C (p=0.048), while diversity and dominance remained comparable across areas (*p*>0.05; **Figure 5D, Supplementary Figure 4E**). Beta diversity analysis showed significant compositional differences (Bray-Curtis R^2^=0.10, *p*=0.044; Jaccard R^2^=0.084, *p*=0.001), although the extent of the differentiation depended on the distance metric used. Pairwise comparisons (Jaccard) discriminated all three zones (**Supplementary Figure 4F**), whereas abundance-based differences (Bray-Curtis) were significant only between Areas A and C (**Figure 5E; Supplementary Table 3**), suggesting that the differences in community composition were driven more strongly by taxon presence/absence than by differences in relative abundance. Consistent with this spatial difference, only 65 genera (20.3%) were shared across all areas (**Supplementary Figure 4G**). Taxonomic profiles also varied among areas. *Fusarium*, *Exophiala*, and *Hanseniaspora* predominating in area A, *Epicoccum* and *Candida* in area B; whereas area C was distinguished by *Rhodotorula*, *Thyridium*, *Epicoccum*, and *Hanseniaspora* (**Figure 5F**). However, ANCOM-BC2 identified no significantly differentially abundant fungal taxa.

Microeukaryotes richness remained comparable across the unit (Observed species *p*>0.05; **Figure 5G**). Nonetheless, beta diversity analysis revealed significant spatial structuring (Jaccard R^2^= 0.098, *p*=0.001), with pairwise comparisons distinguishing all three areas (Jaccard, **Supplementary Table 3**; **Figure 5H**). Reflecting this spatial compartmentalization, the shared core comprised just 50% of genera (**Supplementary Figure 4H**), while area-specific taxa were detected primarily in Areas B (18.2%) and C (13.6%), with none detected exclusively in Area A. Taxonomic composition also varied markedly across areas. *Vermamoeba* reached its highest prevalence in Area A (71%), *Naegleria* was restricted to Area B (25%), while Area C was dominated by *Acanthamoeba* (80%) and showed a higher prevalence in *Colpoda* (53%). (**Figure 5I**). However, high within-group variability limited statistical support, with only five taxa showing marginal evidence of differential prevalence: *Diplophrys*, *Nuclearia*, *Colpoda*, *Vermamoeba* and *Cryptodifflugia* (Fisher’s exact test, BH-adjusted *q*=0.08).

Overall, these results indicate that the three ICU areas harbored distinct bacterial, fungal, and microeukaryotic communities. Although the strength of spatial structuring varied among microbial kingdoms, Area A consistently exhibited the greatest divergence from B and C, suggesting persistent ecological compartmentalization within the ICU water system.

### 3.6. Sink-associated communities shift across operational periods

Comparison of alpha-diversity metrics between the Empty (post-ICU) and Multifunctional periods revealed no significant differences (*p*>0.05; **Figure 6A**, **Supplementary Figure 5A**). However, beta-diversity analysis demonstrated significant bacterial compositional shifts between periods for both Bray-Curtis (R^2^=0.116, *p*=0.001; **Figure 6B**) and Jaccard (R^2^=0.043, *p*=0.001; **Supplementary Figure 5B**) distances. This compositional reorganization occurred despite a substantial shared core, with 971 genera (81.5%) detected in both periods (**Supplementary Figure 5C**). The Empty period was enriched in *Flavobacterium*, *Novispirillum*, and *Comamonas*, whereas *Magnetospirillum*, *Elizabethkingia*, and *Stenotrophomonas* increased during the Multifunctional period (**Figure 6C**). ANCOM-BC2 identified 136 differentially abundant taxa, including 74 enriched in the Empty period and 62 during the Multifunctional period (**Supplementary Table 5**). Furthermore, longitudinal spline analysis (TrendySpliner) (**Figure 6D**) identified significant temporal trajectories for five core genera (*Magnetospirillum*, *Flavobacterium*, *Cloacibacterium*, *Azospira*, *Burkholderia*), while the remaining dominant genera stayed stable over time.

**Figure 6.**
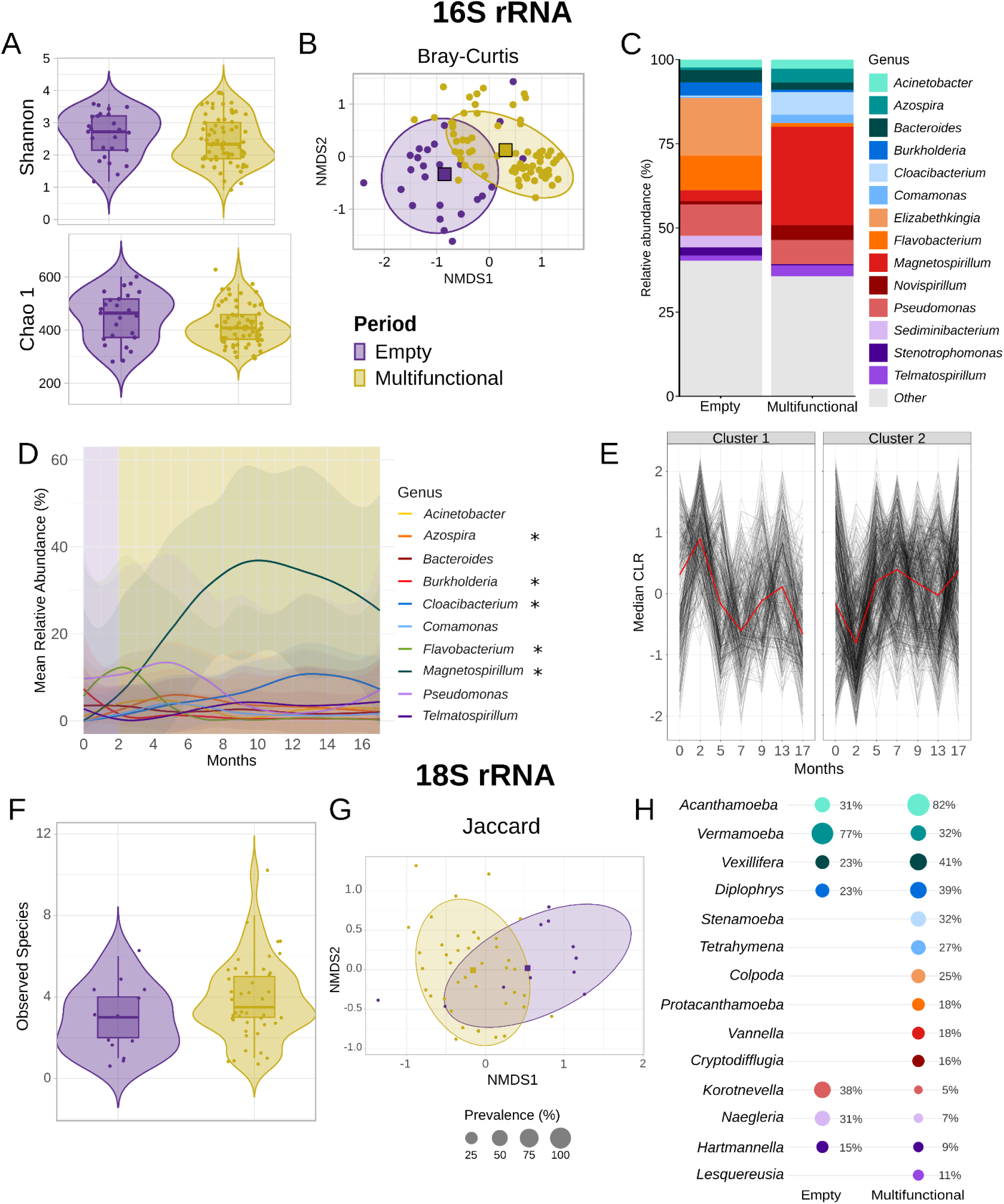
Temporal variations of microbial diversity and composition across periods. Bacterial communities (n=93)**: (A)** alpha-diversity metrics (Shannon index and Chao 1 richness), **(B)** NMDS ordination based on Bray-Curtis dissimilarities; **(C)** relative abundance (%) of the most abundant taxa (Other < 2%); **(D)** temporal trends of the 10 most abundant bacterial genera. Significance (*) indicates a significant non-zero temporal trend based on the “Trendyspliner” implemented in the splinectomeR package. **(E)** Self-Organizing Map (SOM) showing two broad temporal profiles of bacterial taxa, visualized using median CLR-transformed abundance for each cluster. Microeukaryotic communities **(18S rRNA,** n=57)**: (F)** observed richness; **(G)** NMDS plot based on the Jaccard dissimilarity matrix; **(H)** prevalence (%) of the most prevalent detected taxa (n=14).

Self-Organizing Maps (SOMs) further captured the continuous temporal trajectories of bacterial taxa. Across 50 training runs, the maps consistently yielded a quantization error (0.79±0.01) but greater variability in topographic error (0.44±0.024). Silhouette-optimized Ward’s clustering partitioned the map into discrete groups; however, low cross-run stability (mean Adjusted Rand Index = 0.32) demonstrated that these taxa organize along a dynamic continuum rather than inside rigid ecological modules. Treating these partitions strictly as heuristic visual summaries (**Figure 6E**), two inverse median CLR profiles emerged pivoted at the operational shift: Cluster 1 peaked during the Empty period (Months 0–2) before declining, temporarily recovering, and dropping below baseline by Month 17; conversely, Cluster 2 dipped initially before expanding across Months 2–7 and maintaining elevated abundance thereafter.

Microeukaryotic richness remained stable across both operational periods (*p*>0.05; **Figure 6F**). Despite substantial data dispersion, PERMANOVA detected a statistically significant effect of period on community composition based on the Jaccard index (R²=0.08, *p*=0.001; **Figure 6G**), reflected in a restricted shared core of just 31.8% (**Supplementary Figure 5E**), a markedly lower temporal persistence than observed in prokaryotes. Fisher’s exact test revealed no significant differences in prevalence for any genus between the two periods (**Figure 6H**).

## 4. Discussion

This study presents a unique longitudinal view of the multi-kingdom microbial communities inhabiting former ICU sinks during the unit’s transformation into a multifunctional space with entirely different activities. By integrating bacterial, fungal, and microeukaryotic communities with co-occurrence network analysis, we show that sink microbiomes combine a conserved microbial background with distinct spatial signatures and structured associations among microbial groups. Together, these findings reveal an ecosystem shaped by the interplay between ecological persistence, spatial differentiation, and changing environmental conditions.

### 4.1. A conserved multi-kingdom core with sink-specific microbial signatures

Consistent with other surveys of hospital water systems, the ICU bacterial community was dominated by Pseudomonadota and Bacteroidota (Kelly et al., 2023). While a baseline of taxa was shared with other hospital units (29% shared genera), sink microbial communities remained spatially differentiated, indicating that a common microbial background coexists with pronounced local structuring. This combination of ward-specific clustering and a shared baseline is consistent with recent longitudinal culturomics-based observations in hospital sinks, which similarly identified facility-wide inhabitants such as *Pseudomonas* and *Stenotrophomonas* alongside localized variation (Laço et al., 2024).

The persistent bacterial core, representing a mean relative abundance of 77%, defines a highly conserved genus-level ecological niche within former-ICU sinks. While drain-specific literature remains scarce, our cohort is largely supported by the longitudinal ICU study of Chopyk *et al*. (2020), which identified a bacterial core including *Bacillaceae*, *Streptococcus*, *Ralstonia*, and *Herbaspirillum,* all of which were also detected in our study except *Cutibacterium* (Chopyk et al., 2020). As *Cutibacterium* is the most common bacterium of the human skin microbiome (Rozas et al., 2021), its detection may reflect contributions from human-associated biomass rather than persistent drain colonization.

The ubiquitous Pseudomonadota identified here (*Pseudomonas*, *Comamonas*, *Burkholderia*, *Sphingomonas*, and *Sideroxydans*) are frequently reported in hospital sink-drains (Bowie et al., 2025; Healy et al., 2025; Kotay et al., 2026; Laço et al., 2024; Oberauner et al., 2013; Pirzadian et al., 2020). Their persistence is consistent with traits favoring survival in aquatic built environments, including motility, biofilm formation and tolerance to environmental stress (Cooper et al., 2023; de Vries et al., 2019; Fazli et al., 2014; Rasamiravaka et al., 2015). This aligns with the rapid rebound of viable, carbapenem-resistant Pseudomonadota (*Pseudomonas* and *Cupriavidus*, both captured here) following sink disinfection (Bowie et al., 2025). In addition, traits such as xenobiotic degradation and heavy-metal detoxification in *Comamonas* (Ryan et al., 2022) and exopolysaccharide-mediated stress resistance, such as chlorine disinfection in *Sphingomonas*, shield the community against thermal, osmotic, and antimicrobial stresses and may explain the persistence of these core taxa (Banerjee et al., 2021; Sun et al., 2013).

As for the prokaryotic community, a robust fungal core dominated the total relative abundance (81%), driven by Ascomycota and Basidiomycota, which are ubiquitous in environmental and human-associated microbiomes (Dissanayake and Liu, 2025; Nenciarini et al., 2024; Zhang et al., 2024). Core genera, including *Exophiala*, *Fusarium*, and *Malassezia,* replicate patterns from university restroom sinks, supporting their adaptation to anthropogenic water systems (Withey et al., 2023). *Exophiala* may persist through dense extracellular matrices that withstand disinfection (Kirchhoff et al., 2017), while the frequent recovery of *Candida* and *Malassezia* species from hospital sinks is consistent with their established role as persistent sink-associated fungi (Jencson et al., 2017) as a result of anthropogenic impacts in hospital-associated biotic systems (Kumar et al., 2019; Rahimlou et al., 2025). The protist community was dominated by taxa commonly found in drinking water distribution systems (Inkinen et al., 2019), including Ciliophora and diverse ameboid groups belonging to Discosea, Tubulinea, Evosea and Heterolobosea (Delafont et al., 2016). Within the Discosea, *Acanthamoeba*, the unique core protozoan (>70% of prevalence), represents a well-recognized plumbing colonizer previously linked to clinical infections and recovered from hospital taps, dialysis units, and emergency showers (Banerjee et al., 2025; Chomicz et al., 2024).

Despite a shared unit-wide core, individual sinks have distinct microbial signatures. While this sink-specific pattern was strongest and most significant in bacterial populations, the trend extended to both fungal and protozoan communities, demonstrating that the specific microenvironment and history of each sink play a critical role in shaping its local ecosystem. Individual sinks therefore exhibited distinct microbial signatures superimposed on a common ecological background, suggesting that sink communities are structured at multiple spatial scales: broad environmental filtering establishes a persistent community backbone, whereas local conditions and sink history further shape its composition. Similar micro-scale uniqueness has been reported in academic (Cruz et al., 2026) and residential (Hill et al., 2026) plumbing systems, suggesting that localized microbial signatures may be a general property of engineered water environments.

### 4.2. Inter-kingdom microbial networks and pathogen integration in ICU sink reservoirs

Overall, intra-kingdom co-occurrences were significantly more common than inter-kingdom associations, and taxa within the same phyla tend to cluster within the same network groups showing a phylogenetic signal. This pattern is consistent with environmental filtering and shared niche adaptation among related taxa (Emerson and Gillespie, 2008), a phenomenon similarly documented in soil (Siles et al., 2021), plant (Ren et al., 2023), and human microbiomes (Pérez-Cobas et al., 2020a). Both intra- and inter-kingdom exhibited non-random, highly modular topologies, indicating that sink communities are organized into discrete but interconnected ecological modules rather than through neutral and equally probable associations. This high compartmentalization indicates that ICU drains harbor dynamic yet stable core modules governed by a complex interplay of deterministic environmental selection and stochastic community assembly (Zhou and Ning, 2017).

Despite the predominance of within-kingdom associations, the inter-kingdom networks revealed connections that highlight the potential contribution of non-bacteria in the organization and persistence of microbial communities. For instance, among some core fungi taxa, *Candida*, *Malassezia*, and *Rhodotorula* can form mixed bacterial-fungal biofilms (Novak Babič et al., 2020), while free-living amoebae and ciliates act as “Trojan horses,” engulfing and shielding internalized bacteria, such as *Legionella*, *Burkholderia* and *Pseudomonas* (all identified in our study) from biocides, antibiotics, and other environmental stress (Mondino et al., 2020; Morón et al., 2024; Price et al., 2024). Such interactions may provide protected niches that facilitate bacterial survival and dispersal and may contribute to stress and antimicrobial tolerance (Balczun and Scheid, 2017; Morón et al., 2024; Price et al., 2024).

Notably, the co-occurrence networks included several clinically relevant bacterial genera including *Enterobacter*, *Staphylococcus*, and *Streptococcus*, as well as the opportunistic fungus *Aspergillus.* Several representatives of these genera are included in the WHO bacterial or fungal priority pathogen lists, which recognize their public health importance and, for bacterial pathogens, their relevance to antimicrobial resistance (“WHO bacterial priority pathogens list, 2024,” n.d.). Their integration in the network reinforces the role of hospital sinks as persistent reservoirs of healthcare-associated microorganisms (reviewed by McCallum and Hall, 2025), as anchored in our cohort by core opportunistic genera (*Klebsiella*, *Enterobacter*, *Citrobacter*, *Escherichia*, *Serratia*, *Pseudomonas*) and phenotypically confirmed antibiotic resistance across all them, including the carbapenemases VIM or GES previously mapped in this unit (Aracil-Gisbert et al., 2024; “ISPB 2026 | Programme point - Hospital Water Systems Drive the Maintenance and Evolution of Acquired Antimicrobail Resistance via Robust Plasmid and Clonal Networks,” n.d.). Together, these findings link the ecological complexity of sink communities to the presence of clinically relevant and antimicrobial-resistant bacteria.

### 4.3. ICU remodeling and spatial location shape sink microbial community assembly

Microbial communities in former-ICU sinks were spatially structured across the unit’s specific physical areas, with differences in composition and diversity associated with the healthcare-associated activities. Across multiple analyses, Area A consistently showed the greatest divergence from Areas B and C, particularly in bacterial communities, and exhibited higher bacterial richness than both areas. Area A was the most physically distant and, after remodeling, was assigned to office and non-clinical activities, underscoring the role of anthropogenic activity and environmental conditions in shaping the observed spatial microbial structuring. This interpretation aligns with evidence that sink microbiomes are affected by external pressures, including chlorine regimens, faucet hydrodynamics (Bourdin et al., 2026), and disinfection protocols (Dai et al., 2020).

Across our analyses, the primary drivers of multi-kingdom community structure were the ICU operational period and spatial area, followed by sink-level effects, reflecting anthropogenic impacts on local ecology. Rather than manifesting abrupt replacement of microbial communities, the transition from an empty-post ICU to a multifunctional space unfolded along a continuous temporal gradient, driven by inverse taxonomic trajectories. This pattern is consistent with a longitudinal study by Chopyk *et al*., who documented significant shifts in bacterial diversity and composition before, during, and after an ICU renovation (Chopyk et al., 2020). These observations indicate that changes in the physical and functional environment can reshape sink microbial communities without necessarily eliminating their underlying ecological structure.

### 4.4. Limitations

Our findings must be interpreted in the context of two primary methodological constraints. First, statistical power was uneven across kingdoms, as low biomass in the fungal and microeukaryotic cohorts resulted in substantial amplification dropout and smaller sample sizes, which are required to validate the cross-kingdom network topologies. Second, whereas 16S rRNA reference repositories are relatively comprehensive, ITS and 18S rRNA databases remain less complete and more poorly curated. Although mitigated via multi-database consolidation, a higher fraction of non-prokaryotic taxa could not be resolved to the genus level, restricting our ability to fully capture the taxonomic diversity of the fungal and microeukaryotic populations in the sink environment.

## 5. Conclusions

This longitudinal analysis demonstrates that hospital sink drains harbor complex multi-kingdom ecosystems with a conserved core across prokaryotes, fungi, and other microeukaryotes. This core persisted despite prolonged operational vacancy and subsequent changes in the physical and functional environment from an ICU to a multifunctional unit, highlighting the capacity of sink communities to withstand major environmental transitions. This persistence establishes drains as long-term environmental reservoirs for clinically relevant microorganisms and their antimicrobial resistance determinants. While a persistent core baseline exists, community assembly shifts along a continuous temporal gradient shaped by hospital remodeling and changing functional use, with each sink exhibiting a distinct microbial signature shaped by local environmental conditions. Consequently, the sink drain micro-ecosystem represents a significant yet often overlooked challenge for hospital infection control. The resilience of these multi-kingdom communities necessitates more integrated control strategies that account for the persistent, sink-specific nature of these reservoirs. Future efforts should prioritize consideration of plumbing system design and use, alongside ecological monitoring that addresses the multi-kingdom network to effectively mitigate the risk of healthcare-associated infections and evolution of antimicrobial resistance.

## CRediT authorship contribution statement

IL-A: Data curation, Formal analysis, Investigation, Methodology, Software, Visualization, Writing – original draft, Writing – review & editing; NG-P: Formal analysis, Investigation, Methodology, Writing – original draft, Writing – review & editing; SM-R: Methodology, Software, Writing – review & editing; SS-C: Methodology, Writing – review & editing; FA: Validation, Writing – review & editing; AA-I: Validation, Writing – review & editing; MC-S: Resources, Validation, Writing – review & editing; R-P: Resources, Validation, Writing – review & editing; V-FL: Methodology; RC: Resources, Validation, Writing – review & editing; FB: Conceptualization, Validation, Writing – review & editing; TMC: Conceptualization, Funding acquisition, Investigation, Resources, Supervision, Validation, Writing – review & editing; AE-PC: Conceptualization, Investigation, Resources, Supervision, Validation, Writing – original draft, Writing – review & editing.

## Data availability

All sequences have been entered in the European Bioinformatics Institute database under project accession number PRJEB111259.

## Acknowledgements

This work was supported by the European Commission (MISTAR, AC21_2/00041), and the Instituto de Salud Carlos III (ISCIII; PI24/02027), cofinanced by the European Development Regional Fund (A Way to Achieve Europe program, and by grants from Fundación Francisco Soria Melguizo (CC23140547) and Fundación “la Caixa” (grant agreement LCF/PR/HR25/52450012, HR2025-00303). IL-A is supported by the program ‘Ayudas para la contratación de ayudantes de investigación’ of the Comunidad de Madrid (n° PEJ-2024-AI/SAL-GL-31755). NG-P is supported by grants from Fundación Francisco Soria Melguizo (CC23140547) and Fundación “la Caixa” (grant agreement LCF/PR/HR25/52450012, HR2025-00303). SM-R is supported by the Carlos III Health Institute (ISCIII) (n° PI23/01036). AEP-C is supported by the Program ‘Ayudas de atracción de talento investigador César Nombela’ from the Comunidad de Madrid (n° 2023-T1/SAL-GL28953) and by the Carlos III Health Institute (ISCIII) (n° PI23/01036).

