## Supplementary material for "Spatiotemporal dynamics and stability of multi-kingdom microbial communities in hospital sinks acting as persistent pathogen reservoirs": Supplementary_Ambohades_et_al.pdf

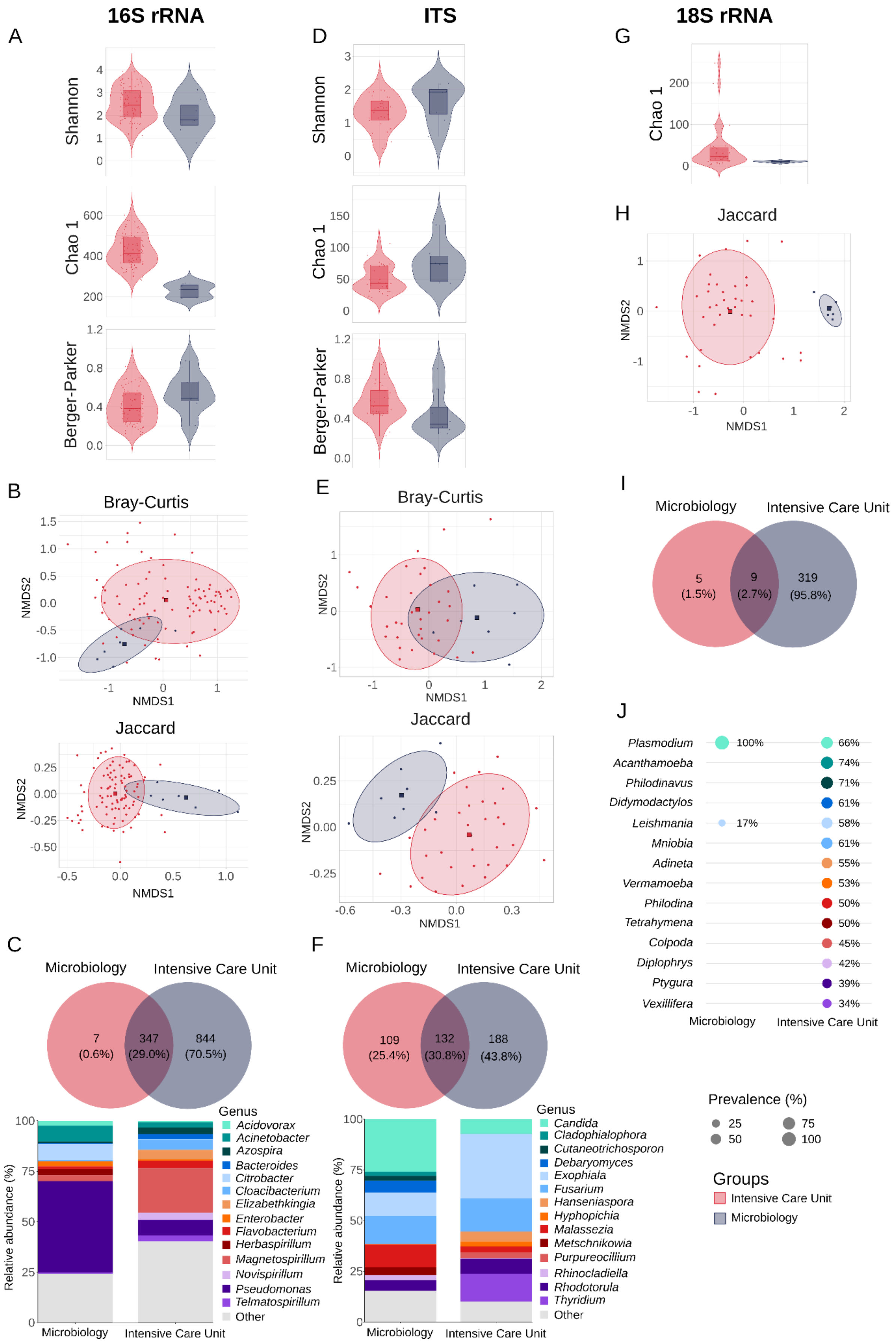
16S rRNA
ITS
18S rRNA
A
D
G
Shannon
Chao 1
Berger-Parker
H
Jaccard
NMDS1
NMDS2
B
E
Bray-Curtis
Jaccard
I
Microbiology
Intensive Care Unit
5 (1.5%)
9 (2.7%)
319 (95.8%)
J
Plasmodium
Acanthamoeba
Philodinavus
Didymodactylos
Leishmania
Mniobia
Adineta
Vermamoeba
Philodina
Tetrahymena
Colpoda
Diplophrys
Ptygura
Vexillifera
100%
17%
66%
74%
71%
61%
58%
61%
55%
53%
50%
50%
45%
42%
39%
34%
Prevalence (%)
25
50
75
100
Groups
Intensive Care Unit
Microbiology
C
F
7 (0.6%)
347 (29.0%)
844 (70.5%)
109 (25.4%)
132 (30.8%)
188 (43.8%)
Relative abundance (%)
Genus
Acidovorax
Acinetobacter
Azospira
Bacteroides
Citrobacter
Cloacibacterium
Elizabethkingia
Enterobacter
Flavobacterium
Herbaspirillum
Magnetospirillum
Novispirillum
Pseudomonas
Telmatospirillum
Other
Candida
Cladophialophora
Cutaneotrichosporon
Debaryomyces
Exophiala
Fusarium
Hanseniaspora
Hyphopichia
Malassezia
Metschnikowia
Purpureocillium
Rhinocladiella
Rhodotorula
Thyridium
Other

**Supplementary Figure 1. Microbial diversity and composition between the ICU and Microbiology units. 16S rRNA** (n=93, n=7)**: (A)** Alpha-diversity metrics, including Shannon index, Chao 1 richness estimator, and Berger-Parker index. **(B)** Sample distribution represented in a non-metric multidimensional scaling (NMDS) plot based on the Bray-Curtis dissimilarity and Jaccard matrices. **(C)** Venn diagram showing shared taxa and their percentage of the total, and relative abundance (%) of the most abundant taxa (Other < 2%). **ITS** (n=34, n=8)**: (D)** Alpha-diversity metrics, including Shannon index, Chao 1 richness estimator, and Berger-Parker index. **(E)** Sample distribution represented in a NMDS plot based on the Bray-Curtis dissimilarity and Jaccard matrices. **(F)** Venn diagram showing shared taxa and their percentage of the total, and relative abundance of the most abundant taxa (Other < 2%). **18S rRNA** (n=57, n=6)**: (G)** Observed species. **(H)** Venn diagram of shared taxa and the percentage of the total. **(I)** Prevalence (%) of the most prevalent taxa (n=14).

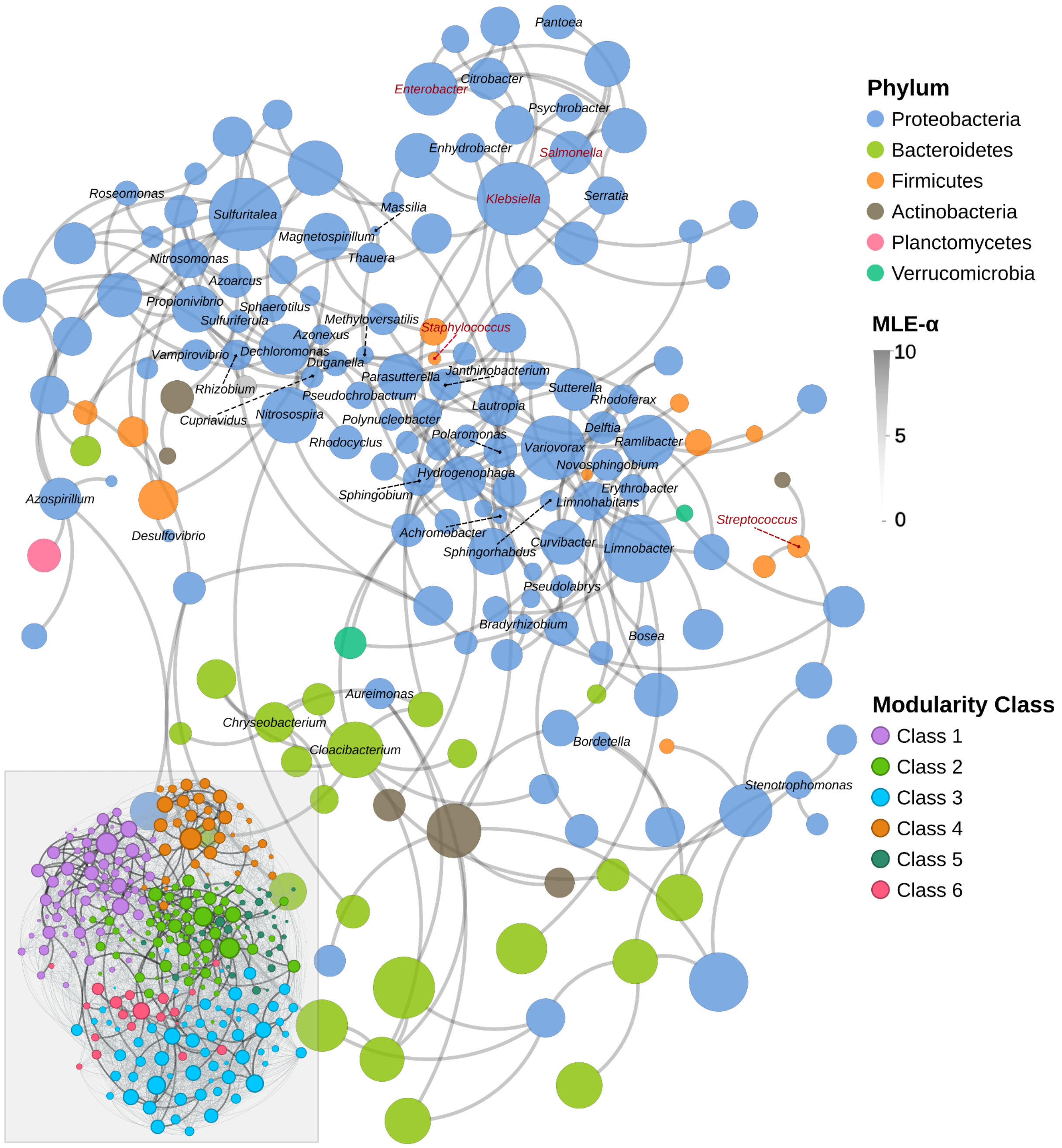


**Supplementary Figure 2. Bacterial intra-domain co-occurrence network.** Phyla are distinguished by color. In the bottom left inset, the network is colored by modularity class (5 clusters): 1 (n=68 taxa), 2 (n=66), 3 (n=63), 4 (n=33), 5 (n=28), 6 (n=19). Only taxa belonging to the core are shown in black. Genera containing species included in the WHO bacterial priority pathogens list are marked in red. The co-occurrence network represents the largest connected component, filtering for significant interactions with MLE-alpha values > 5 and p-values < 0.05. The network colored by modularity showed all significant connections, without filtering by MLE-alpha values, to explore the complete network structure.

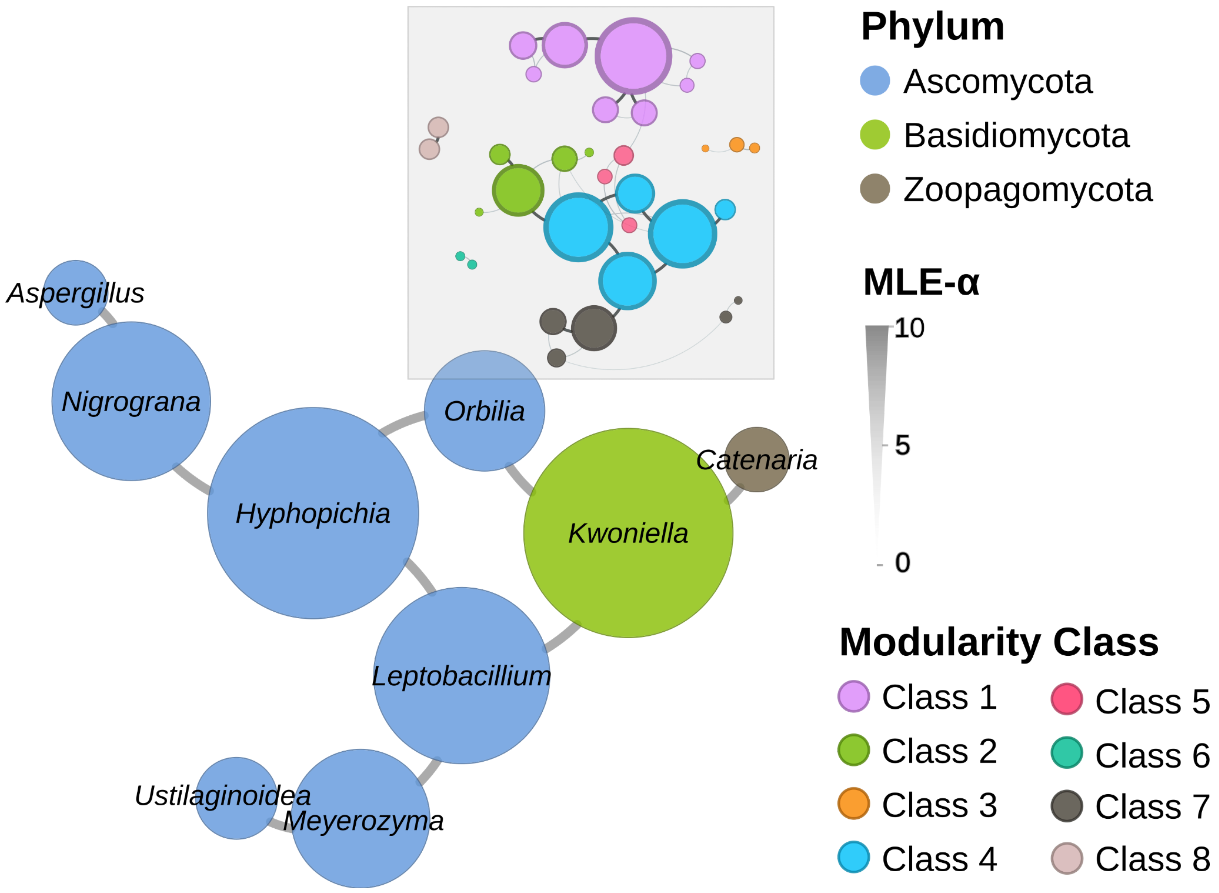


**Supplementary Figure 3. Fungal intra-kingdom co-occurrence network.** Phyla are distinguished by color. In the top center inset, the network is colored by modularity class (5 clusters): 1 (n=8 taxa), 2 (n=5), 3 (n=3), 4 (n=5), 5 (n=3), 6 (n=2), 7 (n=5), 8 (n=2). The co-occurrence network represents the largest connected component exclusively, filtering for significant interactions with MLE-alpha values > 5 and p-values < 0.05. The network colored by modularity showed all significant connections, without filtering by MLE-alpha values, to explore the complete network structure.

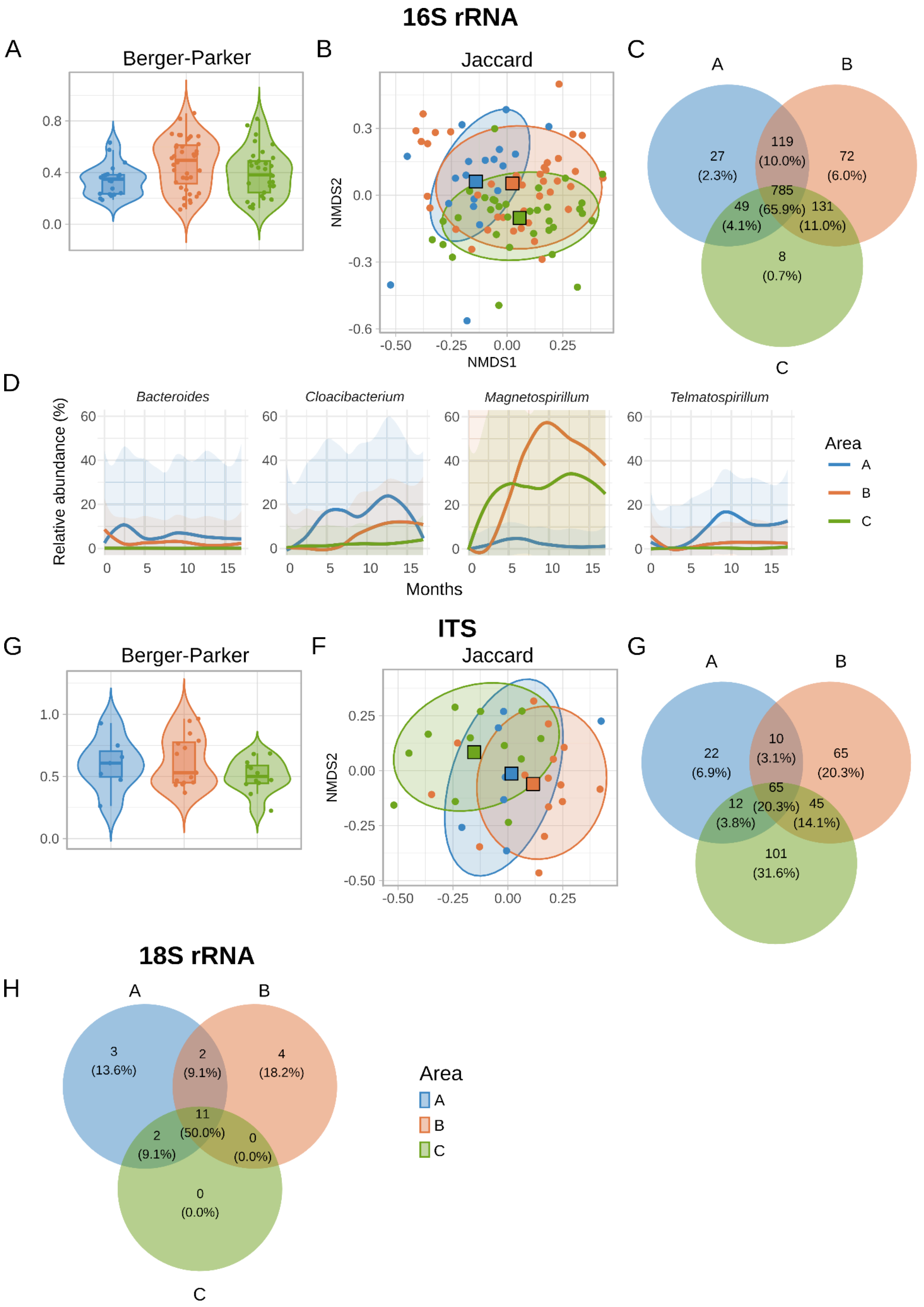


**Supplementary Figure 4. Microbial diversity and composition per area for the 16S rRNA, ITS, and 18S rRNA markers. 16S rRNA** (n=93): **(A)** Berger-Parker index (dominance). **(B)** Samples distribution represented in a non-metric multidimensional scaling (NMDS) plot based on the Jaccard matrix. **(C)** Venn diagram showing shared taxa and their percentage of the total. **ITS** (n=34)**:** **(D)** Temporal trends of taxa exhibiting significant compositional differences across sampling areas over time. Significance was based on the permuSpliner analysis implemented in the splinectomeR package. **ITS** (n=34): **(E)** Berger-Parker index (dominance). **(F)** Samples distribution represented in a non-metric multidimensional scaling (NMDS) plot based on the Jaccard matrix. **(G)** Venn diagram showing shared taxa and the percentage of the total. **18S rRNA** (n=57)**:** **(H)** Venn diagram showing shared taxa and the percentage of the total.

### 16S rRNA

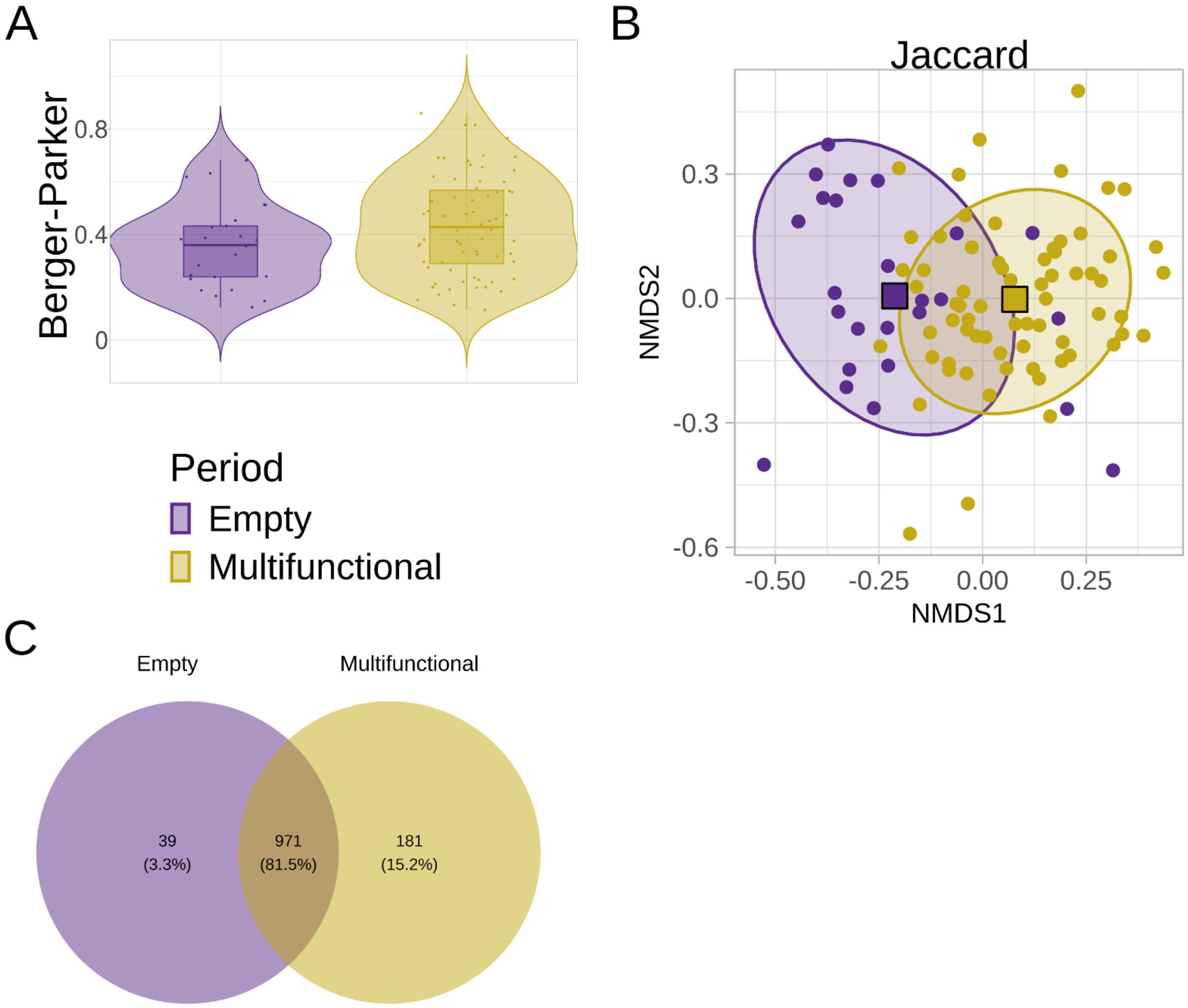


### 18S rRNA

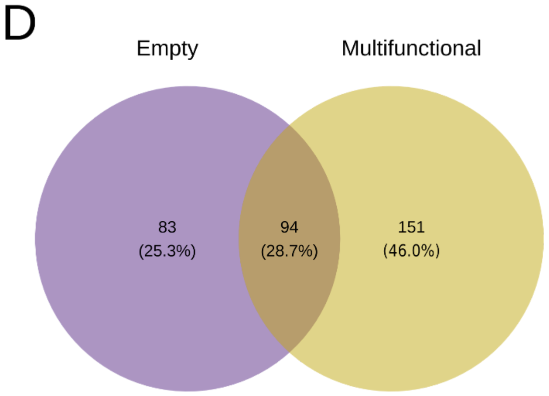


**Supplementary Figure 5. Microbial diversity and composition per period for the 16S rRNA and 18S rRNA markers. 16S rRNA** (n=93)**: (A)** Berger-Parker index (dominance). **(B)** Samples distribution represented in a non-metric multidimensional scaling (NMDS) plot based on the Jaccard matrix. **(C)** Venn diagram showing shared taxa and the percentage of the total. **18S rRNA** (n=57)**: (D)** Venn diagram showing shared taxa and the percentage of the total.
